# BDNF in Ventrolateral Orbitofrontal Cortex to Dorsolateral Striatum Circuit Moderates Alcohol Consumption and Gates Alcohol Habit

**DOI:** 10.1101/2025.01.09.632255

**Authors:** Sowmya Gunasekaran, Jeffrey J. Moffat, Joshua D. Epstein, Khanhky Phamluong, Yann Ehinger, Dorit Ron

## Abstract

BDNF plays a crucial role in shaping the structure and function of neurons. BDNF signaling in the dorsolateral striatum (DLS) is part of an endogenous pathway that protects against the development of alcohol use disorder (AUD). Dysregulation of BDNF levels in the cortex or dysfunction of BDNF/TrkB signaling in the DLS results in the escalation of alcohol drinking and compulsive alcohol use. The major source of BDNF in the striatum is the prefrontal cortex. We identified a small ensemble of BDNF-positive neurons in the ventrolateral orbitofrontal cortex (vlOFC), a region involved in AUD, that extend axonal projections to the DLS. We speculated that BDNF in vlOFC-to-DLS circuit may play a role in limiting alcohol drinking and that heavy alcohol use disrupts this protective pathway. We found that *BDNF* expression is reduced in the vlOFC of male but not female mice after long-term cycles of binge alcohol drinking and withdrawal. We discovered that overexpression of BDNF in vlOFC-to-DLS but not in vlOFC-to-dorsomedial striatum (DMS) or M2 motor cortex-to-DLS circuit reduces alcohol but not sucrose intake and preference. The DLS plays a major role in habitual behaviors. We hypothesized that BDNF in vlOFC-to-DLS circuitry controls alcohol intake by gating habitual alcohol seeking. We found that BDNF over-expression in vlOFC-to-DLS circuit and systemic administration of BDNF receptor TrkB agonist, LM22A-4, biases habitually trained mice towards goal-directed alcohol seeking. Together, our data suggest that BDNF in a small ensemble of vlOFC-to-DLS neurons gates alcohol intake by attenuating habitual alcohol seeking.

## Introduction

The brain-derived neurotrophic factor (BDNF) belongs to the neurotrophins family ^1^. BDNF plays a vital role in neuronal differentiation and maturation ^2^, synaptic plasticity, learning and memory ^3^. BDNF is highly expressed in the adult rodent brain ^4,5^. The majority of BDNF is stored in presynaptic dense core vesicles and is released in an activity-dependent manner upon neuronal depolarization ^6^. BDNF binding to tropomyosin receptor kinase B (TrkB) receptor activates the PI3K/AKT, PLC/PKC and/or ERK1/2 signaling, leading to the activation of transcription ^7^ or translation ^8^. Dysregulation of BDNF signaling has been implicated in psychiatric disorders, such as depression ^9^, schizophrenia ^9^, and addiction ^10^.

Only 10-15% of alcohol users develop AUD ^11^, implying that there are protective mechanisms that prevent the development of the disorder in the majority of the population. Using rodents as a model system, we and others presented data to suggest that BDNF is part of a protective mechanism that gates the development of heavy alcohol use and abuse (reviews ^12-14^). We found that the activation of BDNF/TrkB/ERK1/2 signaling in the DLS, a region involved in habitual behavior ^15^, keeps alcohol intake in moderation ^16-19^.

The prefrontal cortex (PFC) is the major source of BDNF in the striatum ^20-23^, and we and others reported that chronic excessive alcohol consumption leads to an attenuation of *BDNF* expression in corticostriatal regions ^24-26^, and showed that the reduction of the level of *BDNF* message in the medial prefrontal cortex (mPFC) is mediated via microRNA/s ^25,26^ resulting in escalation of alcohol consumption ^25,26^. We further found that escalation of heavy alcohol intake is mediated by the reduction in membranal localization of the TrkB receptor in the DLS ^27^. Finally, we reported that a polymorphism in the BDNF gene that disrupts BDNF release ^28^ results in compulsive excessive intake and alcohol preference over social interaction ^29^ ^30^. Together, these data suggest that BDNF in corticostriatal circuitry/circuitries gates alcohol drinking behaviors, however, AUD-like phenotypes develop in part when BDNF signaling ceases to function.

The cortical regions that release BDNF into the DLS have not been carefully mapped out. Using a combination of transgenic mouse lines together with a viral-mediated retrograde tracing strategy, we characterized BDNF-expressing cortical neurons that project to the DLS. We found that a small ensemble of BDNF-positive neurons in the vlOFC extend axonal projections to the DLS ^23^. The OFC plays a critical role in decision-making, reward-prediction error ^31-33^, reward information ^34^, and stimulus-outcome behaviors ^35-37^. The OFC has also been identified as a critical region in AUD ^38,39^. Specifically, in humans, alcohol dependence reduces white matter and neuronal density in the OFC ^40-42^, and the connectivity between OFC and striatum is altered in abstinent alcoholics ^43^. In rodents, alcohol affects the activity of OFC neurons ^44,45^. Furthermore, OFC lesions or chemogenetic inhibition increases alcohol drinking ^46-48^ and decreases context and cue-induced reinstatement ^49,50^. Together, these data suggest that the OFC is an important target of alcohol. However, whether and if so, how BDNF in vlOFC-to-DLS projecting neurons affects alcohol drinking behaviors is unknown. We report that *BDNF* expression is reduced in the vlOFC of mice following 7 weeks of intermittent access to 20% alcohol in a 2-bottle choice paradigm (IA20%2BC). We further show that overexpression of BDNF in vlOFC-to-DLS circuit limits alcohol intake by reducing habitual alcohol seeking.

## Materials and Methods

Reagents, preparation of solutions, collection of brain samples, real-time PCR, purchasing of viruses, stereotaxic viral infection, confirmation of viral expression, and behavioral paradigms can be found in the supplementary material.

### Animals

Male and female C57BL/6J mice (6-8 weeks) were purchased from Jackson Laboratory and were allowed one week of habituation before experiments began. Mice were individually housed on paper-chip bedding, under a reverse 12-hour light-dark cycle. Temperature and humidity were kept constant at 22 ± 2°C, and relative humidity was maintained at 50 ± 5%. Mice were allowed access to food and tap water *ad libitum*. All animal procedures were approved by the University’s Institutional Animal Care and Use Committee (IACUC) and were conducted in agreement with the Association for Assessment and Accreditation of Laboratory Animal Care.

### Behavioral paradigms

#### Intermittent Access to 20% Alcohol Two-Bottle Choice (IA20%2BC)

IA20%2BC was conducted as previously described ^51^. Briefly, mice were given one bottle of 20% alcohol (v/v) in tap water and one bottle of water for 24 hours on Monday, Wednesday, and Friday, with 24 or 48-hour (weekend) of alcohol withdrawal periods during which mice consumed only water. The placement of water or alcohol bottles was alternated between each session to avoid side preference. Alcohol and water bottles were weighed at the beginning and end of each alcohol drinking session, and alcohol intake (g/kg of body weight), water intake (ml/kg) and total fluid intake (ml/kg) were calculated. Two bottles containing water and alcohol in an empty cage were used to evaluate the spillage. Alcohol preference ratio was calculated by dividing the volume of alcohol consumed to the total volume of fluid intake.

#### Operant alcohol self-administration

Operant alcohol self-administration (OAS) training was performed as described previously ^52^. First, mice underwent 7 weeks IA20%2BC. Mice drinking more than 12.5g/kg were selected for the experiment. OAS training was initiated under a fixed-ratio (FR) 1 schedule, i.e. one lever press resulted in the delivery of one reward, for four 6-hour FR1 sessions followed by four 4-hour FR1 sessions and finally four 2-hour FR1 sessions. All remaining sessions lasted 2 hours. Mice were then trained on a random-interval (RI) schedule, during which rewards were delivered with random delays following active lever presses according to previous studies ^53-55^. Timepoints were pseudo-randomly assigned by the computer program. Animals first underwent 5 sessions on an RI30 schedule (delays averaging 30 seconds after lever press, with intervals ranging from 0-60 seconds). Mice were then subjected to 5 sessions of RI60 training (intervals ranging from 30-90 seconds). Mice were divided into 2 groups with similar numbers of active lever presses (106.41 ± 40.78 and 116.35 ± 58.35), port entries (96.35 ± 45.73 and 72.61± 14.36) and amount of self-administered alcohol (2.55 ± 0.70 and 2.46 ± 0.79 g/kg/2 hr) and BDNF or mCherry control were overexpressed in the vlOFC-to-DLS circuit. Three weeks following surgery, RI30 was resumed for 5 sessions, followed by RI60 until the end of the experiment. The number of active lever presses and reward port entries, as well as the number of reward deliveries, were recorded during each session. LM22A-4 administration: Thirty minutes prior to a degradation session, mice received intraperitoneal (i.p.) administration of saline or LM22A-4 (100 mg/kg) ^29^.

#### Contingency degradation

Contingency degradation was used as previously described ^56,57^ to test the sensitivity of mice to changes in the response-outcome association. The procedure was conducted across nondegraded (ND) and degraded (D) sessions. During the 2-hour D session a lever was extended, but lever presses produced no consequences. In total, 2-3 degradation sessions were performed. During the ND sessions, alcohol deliveries occurred at a rate that was determined based on each animal’s average reward rate during the four RI60 schedules of ND sessions, prior to degradation.

### Statistical Analysis

D’Agostino-Pearson normality test, Shapiro-Wilk normality test and F-test/Levene tests were used to verify the normal distribution of variables and the homogeneity of variance, respectively. Data were analyzed using the appropriate statistical test, including two-tailed unpaired t-test, one-way ANOVA, two-way ANOVA with and without repeated measures followed by post-hoc test as detailed in figure legends. GraphPad Prism 9 was used for statistical analyses. All data are expressed as mean +/- SEM. Significance was set at *p* < 0.05.

## Results

### Excessive alcohol drinking reduces *BDNF* expression in the vlOFC

We previously found that vlOFC neurons expressing BDNF project to the DLS ^23^. Since BDNF signaling in the DLS is a locus for keeping alcohol drinking in moderation ^18,19^, we hypothesized that normal levels of BDNF in the OFC are required to control the development of heavy alcohol use. We further hypothesized that breakdown in BDNF signaling in OFC-to-DLS circuit promotes the escalation of alcohol intake. To address these hypotheses, we first tested whether excessive alcohol drinking alters *BDNF* levels in the vlOFC. Mice underwent 7 weeks of IA20%-2BC paradigm ^52^ (**Figure 1a**). Male mice consumed an average of 14.47± 1g/kg/24 hrs (**Sup. Figure 1a-c; Sup. Table 1**), and female mice drank an average of 19.27±1.1g/kg/24 hrs (**Sup. Figure 1d-f; Sup. Table 1**). Mice were sacrificed 4 hours after the beginning of the last drinking session (“binge”) and 24 hours after the last drinking session (“withdrawal”), and *BDNF* expression was measured. We found that *BDNF* mRNA in the vlOFC was significantly decreased in male mice that were subjected to 7 weeks of IA20%2BC during both binge and withdrawal as compared to mice consuming water only (**Figure 1b**). We further discovered that alcohol-mediated decrease in *BDNF* mRNA levels is localized to the vlOFC since no changes in *BDNF* expression were detected in the medial OFC (mOFC) (**Figure 1c**). As BDNF neurons in M2 motor cortex send dense projections to the DLS ^23^, we examined the level of *BDNF* mRNA in this region in response to alcohol drinking. As shown in **Figure 1d**, *BDNF* mRNA levels in the M2 of male mice were unchanged by alcohol. In contrast, *BDNF* mRNA levels in the vlOFC, mOFC or M2 of female mice were unaltered during binge drinking and after withdrawal from 7 weeks of IA20%2BC **(Figure 1e-g)**. Together, our data suggest that chronic alcohol consumption reduces *BDNF* expression specifically in the vlOFC of male mice. As a result, all subsequent experiments were performed in male mice.

**Figure 1.**
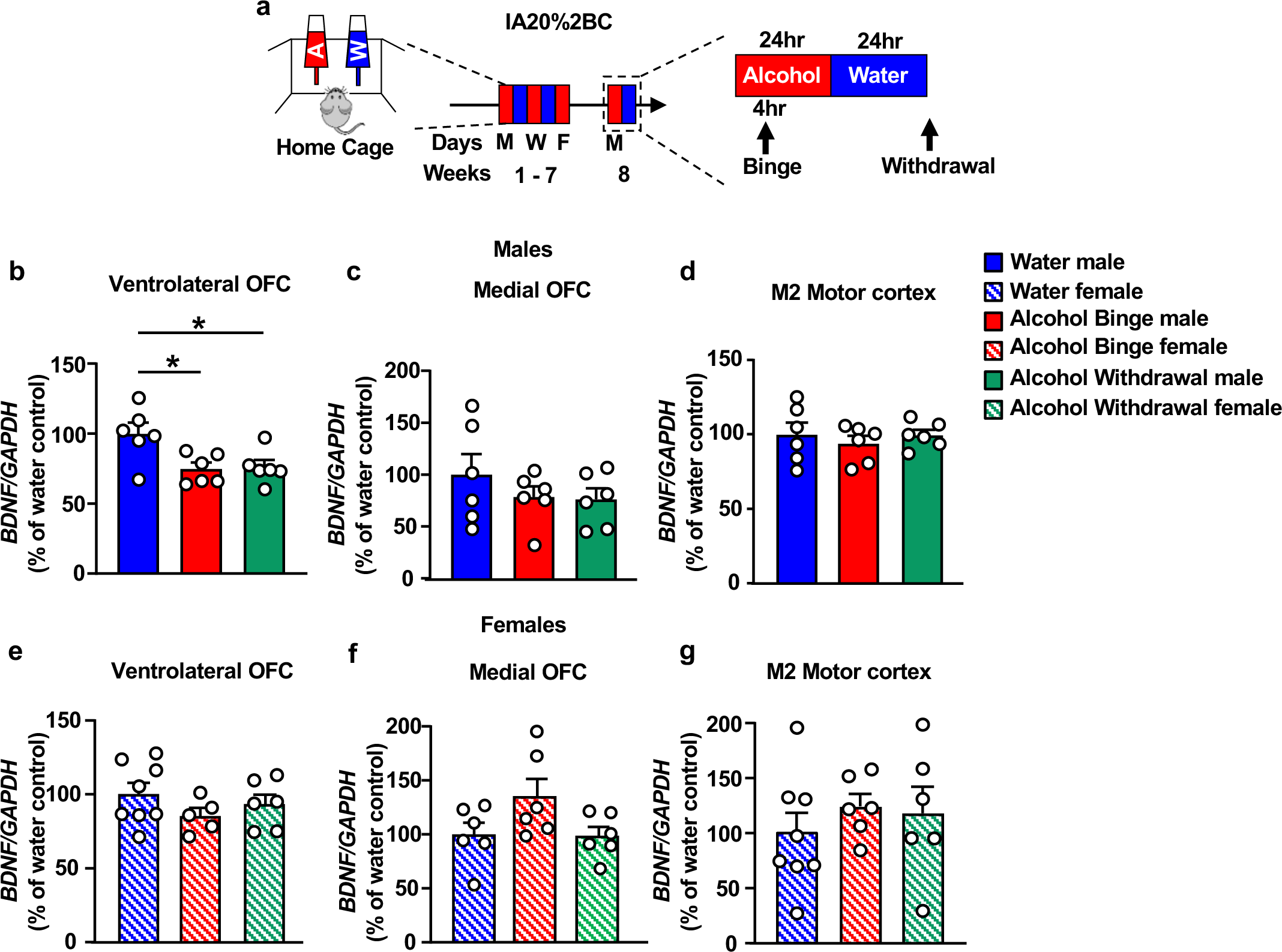
*BDNF* expression is attenuated in the vlOFC but not mOFC and M2 of male but not female mice in response to chronic alcohol drinking and withdrawal. (**a**) Timeline of experiments: Female and male mice underwent 7 weeks of IA20%2BC or water only (**Sup. Table 1**). Four hours after the beginning of the last drinking session (“binge”) and 24 hours after the last drinking session (“withdrawal”), the vlOFC, mOFC and motor cortex M2 were dissected and harvested and the expression of *BDNF* was measured by RT-qPCR. *GAPDH* was used as an internal control. Data are presented as the average ratio of *BDNF* to *GAPDH* ± SEM and are expressed as the percentage of water control. (**b-d**) *BDNF* mRNA in the vlOFC (**b**), mOFC (**c**) and M2 motor cortex (**d**) of male mice (vlOFC, one-way ANOVA: effect of treatment, F _(2,15)_=5.705, p=0.014. Post hoc Dunnett’s multiple comparison test indicates significant differences between water and binge, and water and withdrawal; mOFC, one-way ANOVA: effect of treatment, F_(2,15)_=0.846, p=0.448; M2 motor cortex, effect of treatment, F_(2,15)_=0.354, p=0.707). (**e-g**) *BDNF* mRNA in the vlOFC (**e**), mOFC (**f**) and M2 motor cortex (**g**) of females mice (vlOFC, one-way ANOVA: effect of treatment, F_(2,16)_=1.13, p=0.347; mOFC, effect of treatment, F_(2,15)_=2.88, p=0.087; M2 motor cortex, effect of treatment, F_(2,17)_=0.484, p=0.624). *p < 0.05; ns: non-significant. n=6-8 per group.

### BDNF in vlOFC-to-DLS projecting neurons gates alcohol but not sucrose intake

As described above, excessive alcohol drinking downregulates *BDNF* expression in the vlOFC. If BDNF in vlOFC-to-DLS circuit is gating alcohol intake, then replenishing its levels in this circuitry will revert excessive alcohol intake to moderate levels. To test this possibility, we utilized a circuit-specific strategy to overexpress BDNF in vlOFC neurons that project to the DLS by using the DIO/Cre system enabling *BDNF* expression only in the presence of Cre recombinase (**Figure 2a)** ^23^. AAV2-DIO-BDNF-mCherry virus (1 x 10^12^ gc/ml) was bilaterally infused into the vlOFC and AAVretro-Cre-GFP virus (3 X 10^12^ vg/ml) into the DLS **(Figure 2b-c).** This allowed Cre expression in vlOFC neurons projecting to the DLS visualized by GFP, thus activating Cre-mediated expression of BDNF visualized by mCherry in the vlOFC **(Figure 2c)**. Control animals were bilaterally infected with AAV2-DIO-mCherry (1 x 10^12^ gc/ml) in the vlOFC and AAVretro-Cre-GFP (3 X 10^12^ vg/ml) in the DLS. We found a significant increase of *BDNF* mRNA levels in the vlOFC of animals infected with AAV2-DIO-BDNF-mCherry as compared to animals infected with AAV2-DIO-mCherry **(Figure 2d)**. We further confirmed that *BDNF* in the vlOFC is expressed only in the presence of Cre recombinase (**Sup. Figure 2a-d)**. Three weeks following viral infection enabling maximal BDNF expression, mice were subjected to 7 weeks of IA20%2BC or water only **(Figure 2e)**. We found that overexpression of BDNF in vlOFC to DLS projecting neurons significantly reduced alcohol drinking and preference as compared to control animals **(Figure 2f-g, Sup. Table 1)**. Furthermore, whereas alcohol intake of control mice escalated over time, progressive increase of intake was not detected in mice infected with BDNF in vlOFC neurons projecting to the DLS **(Figure 2f-g)**. Water and total fluid intake were unchanged in the two groups **(Sup. Figure 3a-b)**. Together, these data indicate that escalation of alcohol intake is due in part to the attenuation of BDNF levels in the vlOFC which is rescued by replenishing BDNF in vlOFC-to-DLS projecting neurons. Our data further suggests that BDNF in OFC to DLS circuitry gates escalation of excessive alcohol intake.

**Figure 2.**
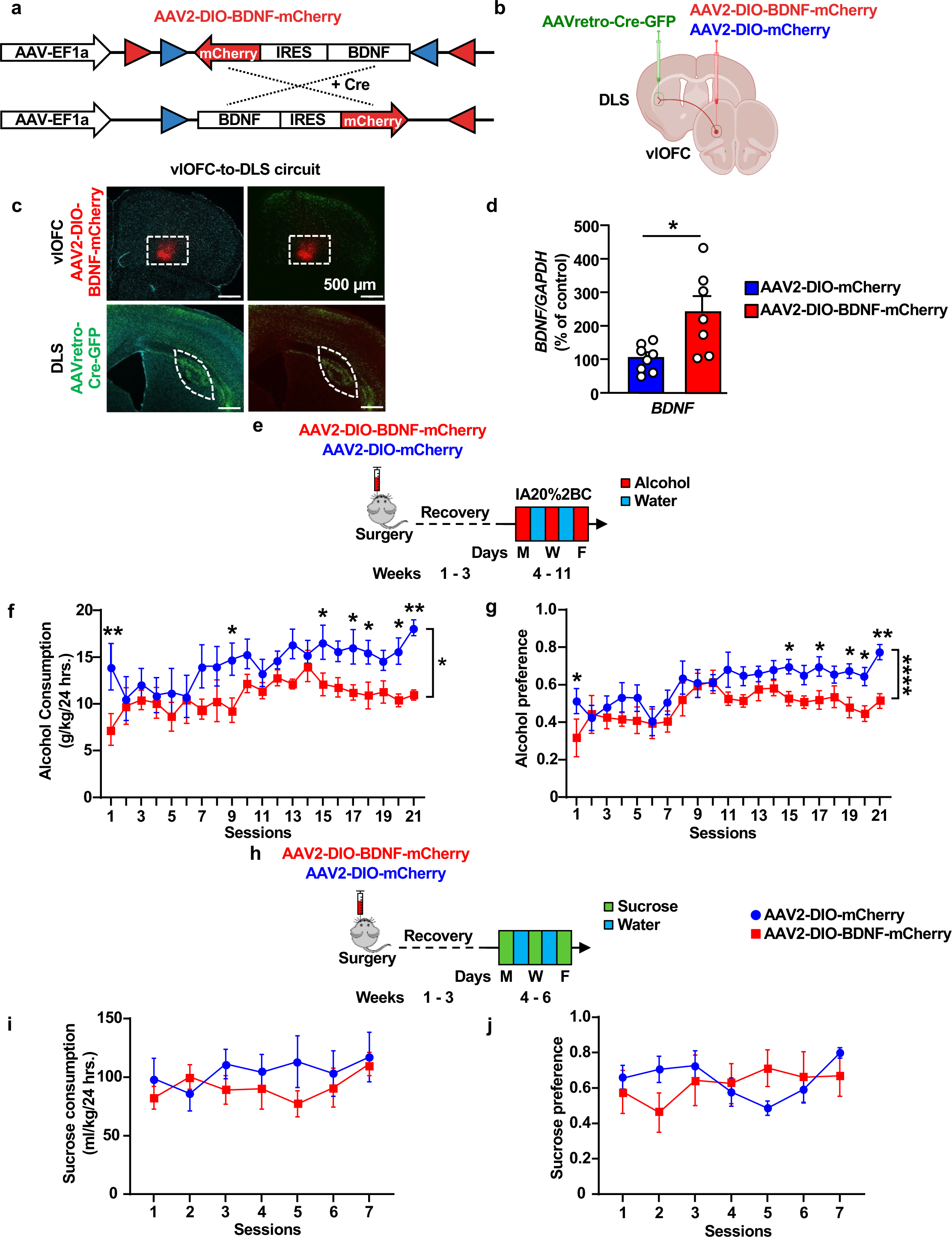
Overexpression of BDNF in vlOFC to DLS circuit moderates alcohol but not sucrose intake. (**a**) Viral strategy of Cre-dependent expression of BDNF: BDNF and mCherry coding sequences are floxed by a pair of loxP (blue triangles) and lox2272 (red triangles) sites. In the absence of Cre recombinase, the BDNF and mCherry coding sequences are inverted relative to the EF1a promoter. When expressed, Cre recombinase inverts the BDNF and mCherry sequences into an active orientation allowing their expression. (**b**) Schematic representation of BDNF overexpression in vlOFC-to-DLS circuit. Mice received bilateral injections of AAV2-DIO-BDNF-mCherry or AAV2-DIO-mCherry in the vlOFC and AAVretro-Cre-GFP in the DLS. (**c**) Representative images depicting targeting of AAV2-DIO-BDNF-mCherry (red) in the vlOFC and AAVretro-Cre-GFP (green) in the DLS. Red cells indicate expression of the Cre and infection with AAV2-DIO-BDNF-mCherry, confirming the overexpression of BDNF in vlOFC neurons projecting to the DLS. (**d**) The vlOFC was dissected 10 weeks after the infusion of AAV2-DIO-BDNF-mCherry or AAV2-DIO-mCherry in the vlOFC and retroAAV-Cre-GFP in DLS and *BDNF* expression was measured by RT-qPCR. *GAPDH* was used as an internal control. Data are presented as the average ratio of *BDNF* to *GAPDH* ± SEM and expressed as the percentage of water control. Unpaired t-test: t _(10)_ = 2.368, p = 0.0394. (**e**) Three weeks after the surgery, mice underwent IA20%2BC for 7 weeks in the home cage (**Sup. Table 1**), and alcohol consumption was recorded. (**f**) alcohol intake (Two-Way ANOVA, effect of BDNF overexpression, F _(1, 264)_ = 47.07, ****p < 0.0001, effect of session, F _(20, 264)_ = 1.791, p=0.02, main effect of interaction, F_(20,264)_=0.714, p=0.810. Post hoc Sidak’s multiple comparison test, between AAV2-DIO-mCherry vs AAV2-DIO-BDNF-mCherry indicates significant difference between conditions on sessions 1,9,15,17,18,20 and 21). (**g**) Alcohol preference was calculated as the ratio of alcohol intake relative to total fluid intake (Two-Way ANOVA, effect of BDNF overexpression, F _(1, 253)_ = 37.90, ****p < 0.0001, effect of session, F _(20, 253)_ = 3.425, *****p < 0.0001, effect of interaction, F _(20, 253)_=0.749, p=0.772. Post hoc Sidak’s multiple comparison test, between AAV2-DIO-mCherry vs AAV2-DIO-BDNF-mCherry indicates significant difference between conditions on sessions 1,15,17,19,20 and 21). (**h**) Mice received a bilateral injection of AAV2-DIO-BDNF-mCherry or AAV2-DIO-mCherry in the vlOFC and AAVretro-Cre-GFP in the DLS and three weeks after the viral injection, mice were subjected to 2-bottle choice with 0.3% sucrose drinking for 7 sessions in the home cage (**Sup. Table 1**). (**i**) Sucrose intake was recorded (Two-Way ANOVA, effect of BDNF overexpression, F _(1, 11)_ = 0.233, p =0.638, effect of session, F _(6, 63)_ = 1.426, p=0.218, effect of interaction, F_(6,63)_=1.605, p=0.160). (**j**) Sucrose preference was calculated as the ratio of sucrose intake relative to total fluid intake (Two-Way ANOVA, effect of BDNF overexpression, F _(1, 11)_ = 0.011, p =0.915, effect of session, F _(6, 58)_ = 1.333, p =0.257, effect of interaction, F _(6, 58)_ = 3.069, p=0.012). Data are represented as mean ± SEM. *p < 0.05, **p < 0.01, ****p<0.0001; n=6-8 per group.

Next, we set out to determine whether BDNF in vlOFC-to-DLS circuit gates the consumption of sucrose, a natural rewarding substance. To do so, a new cohort of animals was subjected to bilateral injections of AAV2-DIO-BDNF-mCherry in the vlOFC and AAVretro-Cre-GFP in the DLS. Control mice were infected with AAV2-DIO-mCherry in the vlOFC and AAVretro-Cre-GFP in the DLS. Three weeks post-viral infusion, mice underwent intermittent access to 0.3% sucrose 2BC paradigm for 2 weeks (**Figure 2h**). We found that *BDNF* overexpression in the vlOFC-to-DLS circuit does not alter sucrose intake and preference compared to control mice **(Figure 2i-j, Sup. Table 1)**. Water consumption and total fluid consumption between the groups were unchanged **(Sup. Figure 3c-d)**. Thus, attenuation of alcohol drinking by BDNF in the vlOFC-to-DLS circuit is not due to changes in palatability and is specific to alcohol. As the striatum plays a role in motor skills ^58^, we investigated whether overexpression of BDNF in vlOFC-to-DLS projecting neurons is due to attenuation of locomotion. As shown in **Sup. Figure 4**, total distance traveled **(Sup. Figure 4b-c)** and velocity **(Sup. Figure 4d)** were similar in the two groups. These results indicate that the behavioral difference in alcohol consumption is not due to changes in motor behavior.

### BDNF in vlOFC-to-DMS or M2-to-DLS projecting neurons does not moderate alcohol intake

A small population of BDNF projecting neurons from the vlOFC also project to the DMS ^23^. Therefore, we investigated whether BDNF in vlOFC-to-DMS projecting neurons modulates alcohol consumption. We used the same circuit-specific viral approach, consisting of a bilateral infusion of AAV2-DIO-BDNF-mCherry in the vlOFC and AAVretro-Cre-GFP in the DMS **(Figure 3a)**, enabling overexpression of BDNF specifically in vlOFC-DMS projecting neurons **(Figure 3b)**. Control mice were infected with AAV2-DIO-mCherry in the vlOFC and AAVretro-Cre-GFP in the DMS. Three weeks post-viral administration, animals underwent IA20%2BC for 7 weeks, and alcohol intake was assessed. We found no significant difference in alcohol **(Figure 3c, Sup. Table 1)**, water and total fluid intake **(Sup. Figure 5a-b),** and alcohol preference (**Figure 3d**) between BDNF-overexpressing mice and control groups. Together, our results suggest that BDNF in vlOFC-to-DLS but not vlOFC-to-DMS circuit moderates alcohol drinking.

**Figure 3.**
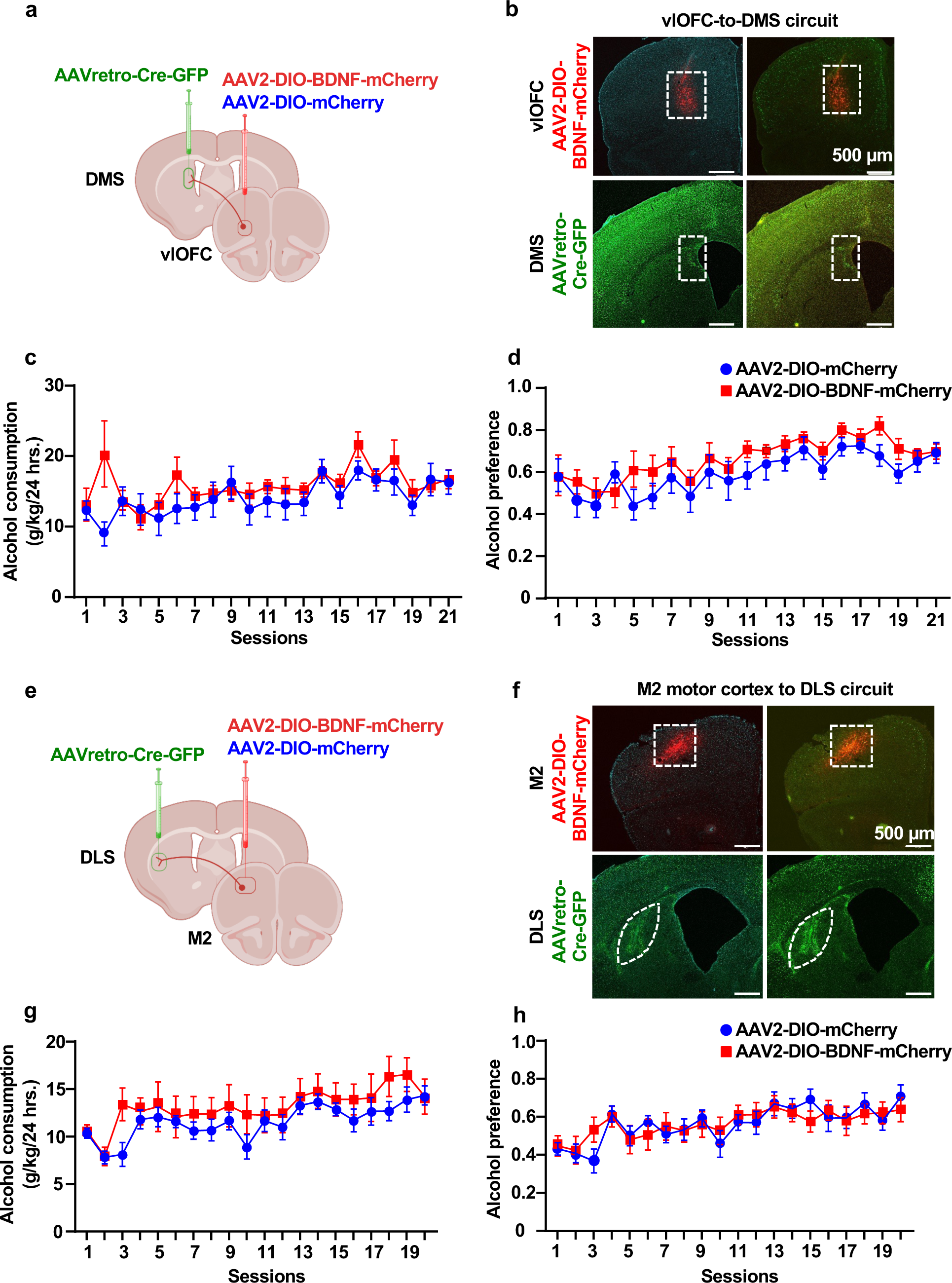
Overexpression of BDNF in vlOFC to DMS or M2 to DLS neurons does not alter alcohol intake. (**a-d**) Overexpression of BDNF in vlOFC to DMS projecting neurons. (**a**) Schematic representation of BDNF overexpression in vlOFC to DMS circuit. Mice received bilateral injections of AAV2-DIO-BDNF-mCherry or AAV2-DIO-mCherry in the vlOFC and AAVretro-Cre-GFP in the DMS. (**b**) Representative image depicting targeting of AAV2-DIO-BDNF-mCherry in vlOFC and AAVretro-Cre-GFP in the DMS. Red cells indicate expression of the Cre and infection with AAV2-DIO-BDNF-mCherry, confirming the overexpression of BDNF in vlOFC neurons projecting to the DMS. (**c**) Three weeks after the surgery, mice underwent IA20%2BC for 7 weeks in the home cage (**Sup. Table 1**), and alcohol/water intake was recorded (Two-Way ANOVA, effect of BDNF overex-pression, F _(1, 17)_ = 1.033, p =0.323, effect of session, F _(20, 330)_ = 3.750, ****p <0.0001, effect of interaction, F _(20,330)_ =1.592, p=0.052. (**d**) Alcohol preference was calculated as the ratio of alcohol intake relative to total fluid intake (Two-Way ANOVA, effect of BDNF overexpression, F _(1, 16)_ = 1.626, p =0.220, effect of session, F _(20, 307)_ = 6.648, ****p < 0.0001, effect of interaction, F_(20,307)_=0.5, p=0.958. (**e-h**) Overexpression of BDNF in M2 to DLS projecting neurons (**e**) Schematic representation of BDNF overexpression in M2 to DLS circuit. Mice received bilateral injections of AAV2-DIO-BDNF-mCherry or AAV2-DIO-mCherry in M2 and AAVretro-Cre-GFP in the DLS. (**f**) Representative image depicting targeting of AAV2-DIO-BDNF-mCherry in M2 and AAVretro-Cre-GFP in the DLS. Red cells indicate expression of the Cre and infection with AAV2-DIO-BDNF-mCherry confirming the overexpression of BDNF in M2 neurons projecting to the DLS. (**g**) Three weeks after the surgery, mice underwent IA20%2BC for 7 weeks in the home cage (**Sup. Table 1**), and alcohol/water intake was recorded (Two-Way ANOVA, effect of BDNF overexpression, F _(1, 16)_ = 0.8408, p =0.372, effect of session, F _(19, 294)_ = 5.445, ****p <0.0001, effect of interaction, F_(19,294)_=0.709, p=0.809). (**h**) Alcohol preference was calculated as the ratio of alcohol intake relative to total fluid intake (Two-Way ANOVA, effect of BDNF overexpression, F _(1, 16)_ = 0.0001175, p =0.991, effect of session, F _(19, 299)_ = 5.541, effect of interaction, F_(19,299)_=0.918, p=0.560). Data are represented as mean ± SEM, n=8-10 per group.

BDNF-expressing neurons in M2, a region essential for motor learning and behaviors ^59^, extend dense projections to the DLS ^23^. We, therefore, assessed whether BDNF in M2-to-DLS projecting neurons alters alcohol consumption. To do so, AAV2-DIO-BDNF-mCherry was infused into the M2 and AAVretro-Cre-GFP into the DLS **(Figure 3e-f)**. Control mice were infected with AAV2-DIO-mCherry in the M2 and AAVretro-Cre-GFP in the DLS. We found that BDNF overexpression in M2 neurons that project to the DLS does not alter alcohol intake **(Figure 3g, Sup. Table 1)** and preference **(Figure 3h)**. Water and total fluid consumption were unchanged between the groups **(Sup. Figure 5c-d)**. These data indicate that, unlike BDNF in vlOFC-to-DLS circuitry, BDNF in neurons that project from M2-to-DLS do not contribute to mechanisms regulating alcohol drinking.

### BDNF in vlOFC-to-DLS projecting neurons protects against the development of habitual alcohol seeking

Next, we set out to elucidate the mechanism by which BDNF in vlOFC-to-DLS circuit gates alcohol intake. The OFC and DLS have been implicated in habit formation and habitual drug seeking ^55,57,60-63^. Alcohol seeking becomes habitual over time ^55,64^. We hypothesized that the downregulation of BDNF level in vlOFC neurons projecting to the DLS during escalation of heavy alcohol intake contributes to the formation of habitual alcohol seeking, which is restored upon replenishing the growth factor in this circuitry. To test the hypothesis, we utilized a contingency degradation procedure to examine habitual alcohol seeking ^52^. Mice first underwent IA20%2BC for 7 weeks (**Sup. Figure 6a-d, Sup. Table 1**) and were then trained to operantly self-administer alcohol by lever pressing for 20% alcohol using a random interval (RI) training of reinforcement (**Timeline, Figure 4a**) which biases lever responding toward habitual actions ^53-55^. Specifically, mice were initially trained on an FR1 schedule for 14 sessions followed by an RI schedule for 5 sessions. Mice were then trained on an RI60 schedule for 5 sessions. Alcohol lever presses were similar in mice that were preassigned to be infected with AAV2-DIO-BDNF-mCherry or AAV2-DIO-mCherry in the vlOFC and AAVretro-Cre-GFP in the DLS (**Suppl. Figure 6e**). Following the initial training, mice were bilaterally infused with AAV2-DIO-BDNF-mCherry or AAV2-DIO-mCherry in the vlOFC and AAVretro-Cre-GFP in the DLS (**Figure 4a**). Three weeks after the surgery, mice underwent 5 additional sessions of RI30 training followed by 5 sessions of RI60 training. We detected fewer active lever presses and rewards in mice overexpressing BDNF in vlOFC-to-DLS circuit compared to the control mCherry mice (**Figure 4b-c**), suggesting that BDNF overexpression in vlOFC-to-DLS circuit contributes to alcohol seeking. Mice trained to self-administer alcohol in a goal-directed manner are sensitive to contingency degradation, while habitually trained mice are not ^53-55^. To determine whether overexpression of BDNF in vlOFC-to-DLS circuit reduces alcohol seeking due to attenuation of goal-directed or habitual alcohol seeking, we conducted a contingency degradation procedure and tested whether habitually trained mice continue to press a lever previously paired with alcohol reward delivery (**Figure 4d**). During degradation sessions, mice received alcohol at a rate equal to their average reward rate during the 4 non-degradation self-administration sessions immediately preceding degradation. As shown in **Figure 4e**, the control AAV2-DIO-mCherry infected mice pressed similarly during nondegraded and degraded sessions, indicating habitual responding. However, AAV2-DIO-BDNF-mCherry-infected mice significantly reduced active lever pressing during the degraded sessions compared to non-degraded sessions, suggesting that these mice are sensitive to degradation of action-outcome contingency **(Figure 4e)**. Port entries during the non-degraded and degraded sessions were unchanged **(Figure 4f).** Finally, using the open field test, we detected no change in locomotion between the groups **(Figure 4g)**. Together, our data suggest that BDNF in the vlOFC-to-DLS circuit reduces habitual alcohol seeking.

**Figure 4.**
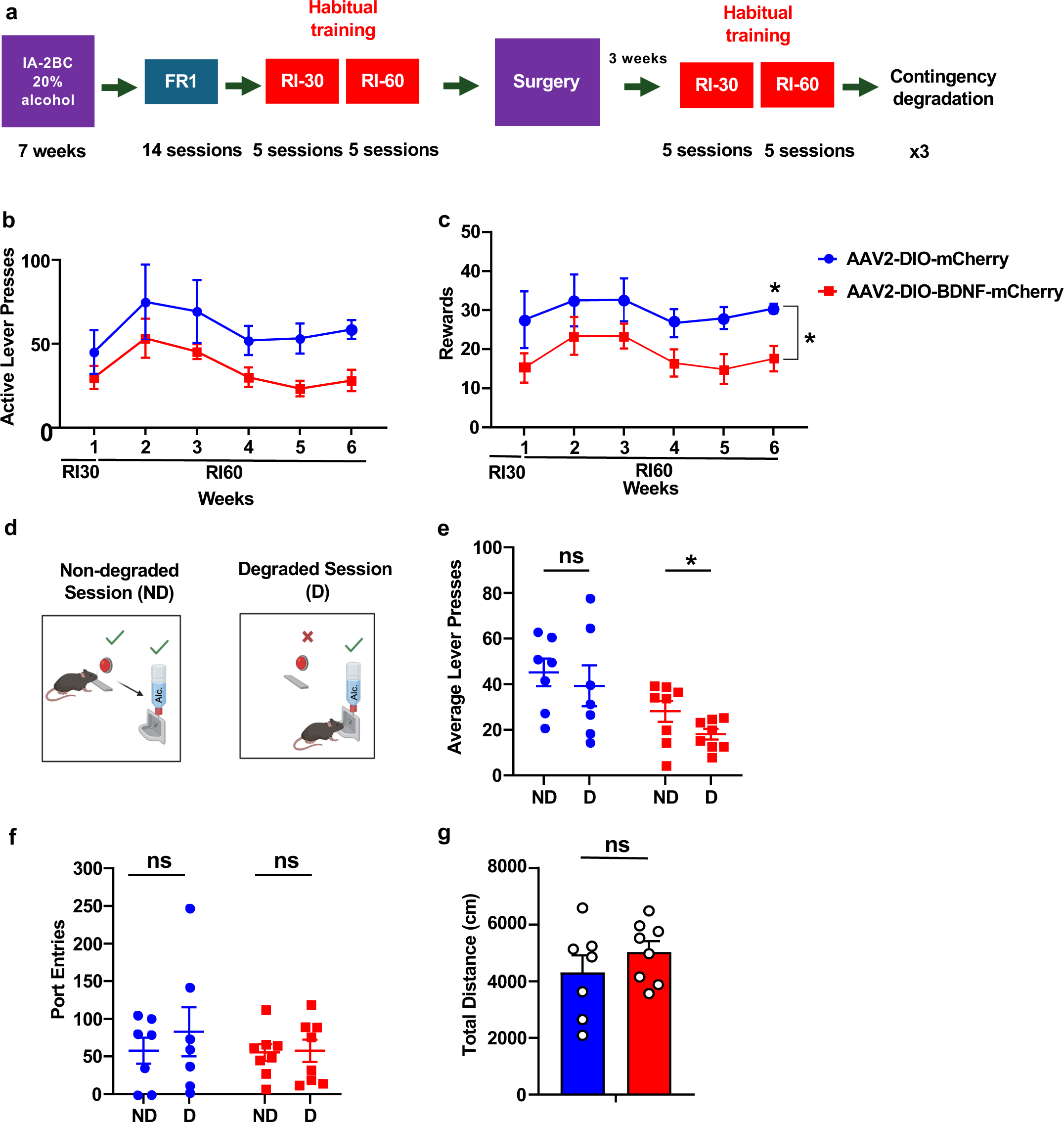
BDNF in vlOFC-to-DLS circuit biases habitual trained mice to goal-directed alcohol seeking. **(a)** Experimental timeline: mice underwent 7 weeks of IA20%2BC alcohol in their home cage (**Sup. Table 1**). Mice were then trained to operant self-administer 20% alcohol during four weeks of FR1 sessions followed by RI training schedule RI30 and RI60 for two weeks and were pseudo-randomly assigned to two groups. One group received bilateral infusions of AAV2-DIO-mCherry in the vlOFC and AAVretro-Cre-GFP in the DLS. The second group received AAV2-DIO-BDNF-mCherry in the vlOFC and AAVretro-Cre-GFP in the DLS. Three weeks after surgery, RI training was resumed. Following a stable RI60 responding, mice underwent contingency degradation testing depicted in (**d**). (**b**) Average of active lever presses during the different sessions after the surgery (Two-way RM ANOVA: Effect of virus F_(1,12)_ = 4.714, p = 0.050; Effect of session F_(1.887, 22.64)_ = 3.138, p = 0.0653; Effect of interaction F_(5,60)_ = 0.205, p = 0.958. Post hoc Sidak’s multiple comparison test indicates a significant difference between conditions in session 6). (**c**) Rewards received during the operant training sessions after surgery (Two-way RM ANOVA: Effect of virus F_(1,13)_ = 6.254, p = 0.026; Effect of session F_(2.162,28.11)_ = 1.879, p = 0.169; Effect of interaction F_(5,65)_ = 0.150, p = 0.979. Post hoc Sidak’s multiple comparison test indicates a significant difference between conditions in session 6). (**d**) The contingency degradation test consists of two types of sessions: nondegraded (ND) and degraded (D). During ND sessions, rewards were delivered on an RI schedule. During D sessions, rewards were delivered at a rate equal to the average of the last week of training, but active lever presses had no effect. The test consisted of an ND session followed by a D session, repeated three times. (**e**) Average active lever presses during ND and D testing sessions in RI schedule of reinforcement (Two-way RM ANOVA: Effect of virus F_(1,13)_ = 6.669, p = 0.022; Effect of session F_(1,13)_ =5.539, p = 0.035; Effect of interaction F_(1,13)_ = 0.367, p = 0.554) (**f**) Average port entries during ND and D testing sessions (Two-way RM ANOVA: Effect of virus F_(1,13)_ = 0.404, p = 536; Effect of session F_(1,13)_ =5.590, p = 0.456; Effect of interaction F_(1,13)_ = 0.416, p = 0.529). (**g**) Open field locomotion test. Mice were placed in an open field and total distance was recorded for 10 minutes (Unpaired t-test: t _(13)_ = 1.044, p = 0.315). Data represented as mean ± SEM. *p < 0.05. ns: non-significant. n=7-8 per group.

### Systemic administration of a TrkB agonist reverses habitual alcohol seeking

Finally, to examine the translational utility of the findings, we utilized the TrkB agonist, LM22A-4 which selectively binds and activates TrkB ^65-68^ (**Figure 5a)** and tested its ability to suppress habitual alcohol seeking, following 7 weeks of IA20%2BC, a new cohort of mice was trained to self-administer 20% alcohol in operant chambers. After 7 initial training sessions on an FR1 schedule, mice began operant training using an RI schedule ^53-55,57^ (**Timeline, Figure 5b, Sup. Table 1**). Mice were subjected to 5 sessions of RI30 and RI60 training followed by contingency degradation (**Timeline**, **Figure 5b**). Mice received an intraperitoneal (i.p.) administration of saline or a TrkB agonist, LM22A-4 (100 mg/kg) ^29^, 30 minutes prior to the degradation session, in a counterbalanced manner. As shown in **Figure 5c**, RI-trained mice treated with saline showed no significant differences in lever pressing between non-degraded and degraded sessions. However, RI-trained mice treated with LM22A-4 exhibited a reduction in lever presses during contingency degradation, compared with non-degraded sessions (**Figure 5c**). Taken together, these data show that treatment with a TrkB agonist biases mice trained to habitually selfadminister alcohol toward goal-directed action selection strategy. Furthermore, these results indicate the potential of utilizing a TrkB agonist to prevent habitual drug seeking.

**Figure 5.**
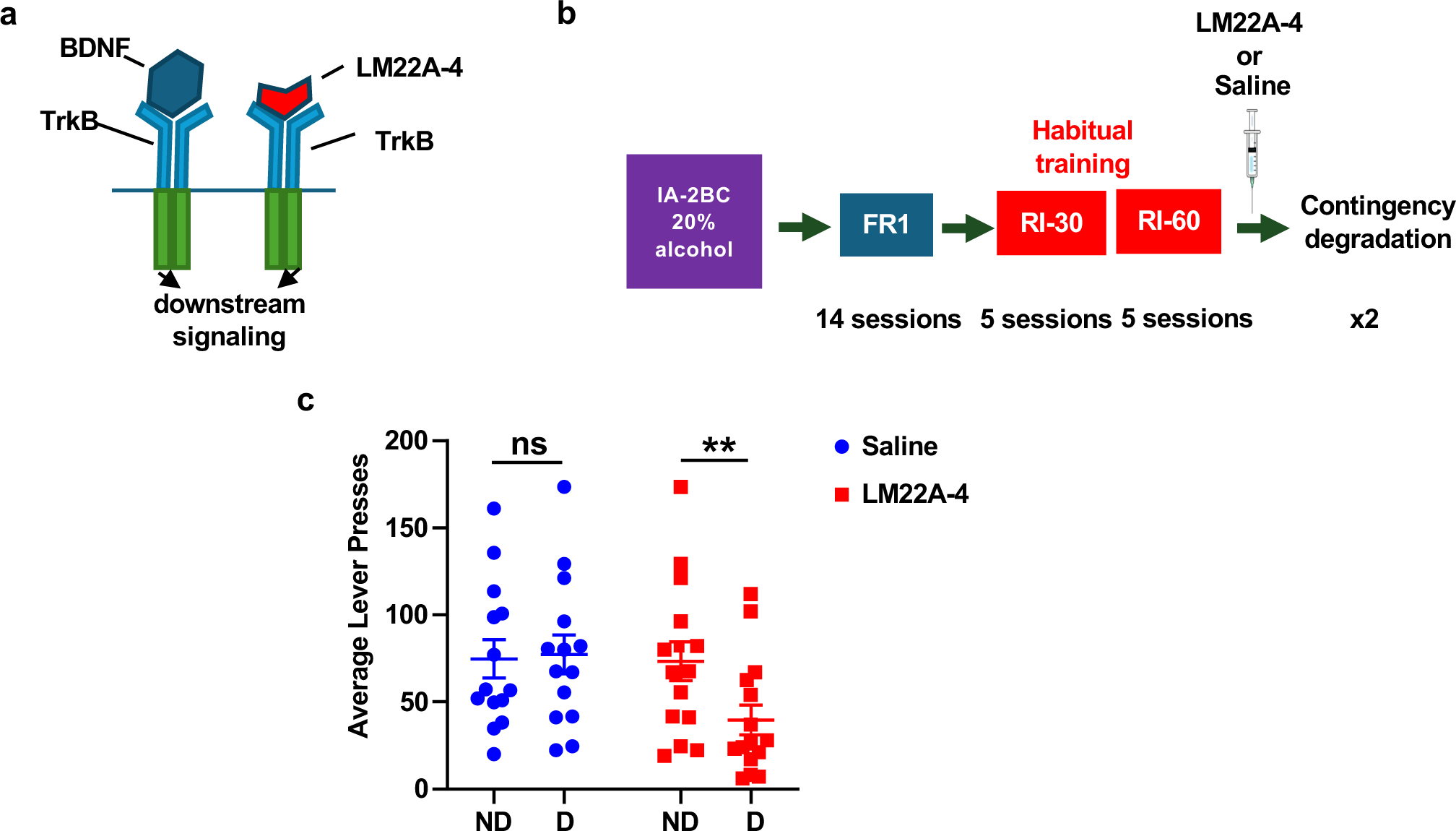
Systemic administration of TrkB agonist LM22A-4 biases habitually trained mice towards goaldirected alcohol seeking. (**a**) LM22A-4 mechanism of action: LM22A-4 binds to and activates TrkB leading to the activation of downstream signaling pathways ^65^. (**b**) Timeline of operant self-administration training. Following 7 weeks of IA2BC-20% alcohol in the home cage (**Sup. Table 1**), mice began operant training for alcohol (20%) self-administration. The first 4 weeks, Monday through Friday, were dedicated to training on a progressive RI schedule with contingency degradation sessions on the final 2 Fridays. Mice were pseudo-randomly assigned and received an i.p. administration of saline or LM22A-4 (100 mg/kg) 30 minutes prior to the first contingency degradation. (**c**) Average lever presses during ND and D testing sessions in RI schedule of reinforcement (Two-Way RM ANOVA, main effect of degradation F _(1, 26)_ = 13.52, p =0.0011, effect of treatment F_(1,28)_=0.93, p=0.342, effect of interaction F_(1,26)_=16.08, p=0.0005). Data are represented as mean ± SEM; **p < 0.01, ns: non-significant. n = 14.

## Discussion

Here, we investigated the potential role of BDNF in vlOFC to circuit in alcohol consumption and habit. We found that chronic excessive alcohol intake decreases *BDNF* levels in the vlOFC but not in mOFC or M2 of male but not female mice. We further discovered that BDNF in a small ensemble of vlOFC neurons projecting to the DLS gates alcohol but not sucrose drinking and habitual alcohol seeking. Together our results suggest that BDNF/TrkB signaling in the vlOFC-DLS keeps alcohol drinking in moderation in part via gating habitual alcohol seeking. Finally, we showed that systemic administration of a TrkB agonist biases habitual alcohol seeking to goal-directed behavior suggesting a utility of TrkB agonist to dampen alcohol intake and habit.

The rodent OFC is a complex brain region containing several functionally distinct subregions, with different neuronal populations extending axonal projections throughout the brain ^35^. We recently showed that the vlOFC sends projections to the DLS ^23^. In rodents, primates, and humans, the vlOFC is an anatomically separate subregion, with distinct corticostriatal circuits linked to specific behavioral functions ^35,69^. We found a significant decrease in *BDNF* mRNA levels in the vlOFC after excessive alcohol drinking in male mice. The mechanism responsible for the attenuation of *BDNF* levels in the vlOFC following long-term binge drinking and withdrawal remains unknown. *BDNF* mRNA is regulated in part by the microRNA (miR) machinery which promotes mRNA degradation or translation repression ^70^. We and others previously showed that repeated cycles of excessive alcohol intake or alcohol vapor exposure and withdrawal decrease *BDNF* expression in the medial PFC (mPFC) of mice and rats ^25,26^ which correlates with increased levels of miR-30a-5p and miR-206 both targeting *BDNF* mRNA ^25,26^. Overexpression of these miRs in the mPFC results in an escalation of alcohol intake while their inhibition reduces excessive drinking ^25,26^. These studies were conducted on male mice; however, evidence suggests sex-specific patterns of miR expression in response to various stimuli ^71,72^. Therefore, it would be of interest to identify the BDNF-targeting miRs increased in the vlOFC after excessive alcohol intake in male vs. female mice, which could be one of the mechanisms underlying the sexual dimorphism we identified in this study.

We found that BDNF overexpression in vlOFC-to-DLS projecting neurons decreases alcohol drinking and habitual alcohol seeking. The OFC is involved in motivation ^35,53^ and modifying stimulus-outcome associations ^35,53,56,73,74^ along with reward seeking ^35,53^. Gourley and colleagues reported that BDNF in the vlOFC is involved in goal-directed decision-making ^56,75^. Specifically, the authors showed that the knockdown of BDNF in OFC interferes with stimulus-outcome and response-outcome associations and that systemic administration of BDNF agonists rescues action-selection associations ^76^. In addition, Pitts and colleagues found BDNF to be involved in the balance between action and habit ^77^ by showing that overexpressing truncated inactive TrkB in vlOFC impedes goal-directed action ^77^. These findings, together with ours, suggest that BDNF in vlOFC-to-DLS circuit biases animals towards goal-directed alcohol seeking and that BDNF in vlOFC-to-DLS circuit gates alcohol intake by altering decision-making.

The DLS has been linked with habitual seeking of drugs and natural rewards in rodents ^78,79,77,80^. To our knowledge, this is the first report to suggest of DLS mechanism that gates habit. It would be interesting to determine whether disruption of DLS signaling in psychiatric disorders resulting in maladaptive habitual and compulsive behaviors can be restored by activation of BDNF signaling in the DLS. In line with this possibility, Pitts et al. reported that inhibition of TrkB signaling in the DLS is sufficient to prevent the development and action of habitual behavior ^77^.

Axonal release of BDNF activates TrkB receptors in the target region. We showed that BDNF-positive neurons from the vlOFC make synapses with DLS neurons ^23^, suggesting that TrkB signaling in the DLS is activated by overexpression of BDNF in vlOFC-to-DLS neurons. TrkB activation promotes the activation of ERK1/2, PLC/PKC or PI3K/AKT signaling ^81,6^. We previously found that BDNF-mediated activation of TrkB in the DLS gates alcohol intake in a mechanism that depends on ERK1/2 but not PI3K/AKT or PLC/PKC signaling ^19^ suggesting that BDNF from vlOFC neurons promotes the activation of TrkB/ERK1/2 leading to transcriptional or translational modifications that in turn gates alcohol intake and habit.

The results herein further highlight the importance of the vlOFC to DLS projection in regulating alcohol drinking. The majority of DLS neurons are dopamine D1 receptors (D1) or dopamine D2 receptors (D2)-expressing medium spiny neurons (MSN) ^82^. Both D1 and D2 MSN express TrkB receptors ^83^. Within the DLS, it is plausible that BDNF signaling via TrkB is differentially regulated depending on the target cell i.e. D1 or D2 MSNs ^83,84,85^. Work from Nestler and colleagues suggests that activation of BDNF-TrkB signaling in D1 versus D2 MSN in the nucleus accumbens generates opposite effects on cocaine and morphine-dependent rewarding behaviors ^83,84^. Further studies are required to gain insight into BDNF-TrkB signaling in behavioral responses to alcohol in subpopulations of DLS neurons.

The vlOFC extends projections to several other brain regions, including the hippocampus, substantia nigra, ventral tegmental area, and several cortical subregions ^86,87^. In addition, work from Gourley and colleagues suggest that the vlOFC projections to the ventrolateral striatum regulates goal-directed food and cocaine seeking ^56^ and Saunders and colleagues recently reported a role of the vlOFC to basolateral amygdala circuit is critical during reward seeking ^88^. Thus, we cannot exclude the possibility that BDNF originating from the vlOFC activates TrkB signaling in one or more of these brain regions and may also play a role in alcohol intake.

We previously reported that administration of LM22A-4 reduced compulsive drinking in the Met68BDNF mutant mice which exhibited compulsive alcohol drinking ^29^. We show herein that LM22A-4 reverses habitual alcohol seeking. LM22A-4 has been shown to have promising effects in preclinical studies for the treatment of several neurological diseases ^89-92^. Together, these data give rise to the potential use of LM22A-4 in AUD.

Overall, these results highlight the importance of a small vlOFC-to-DLS ensemble in regulating alcohol intake. Our data also provide insight into mechanisms that can protect against escalating alcohol consumption.

## Acknowledgements

This study was funded by NIH/NIAAA R37 AA016848 (DR) and NIH/NIAAA R01 AA031832 (DR). BioRender.com was used to create some of the figures.

## Author Contributions

DR conceived the project. DR and YE provided oversight and guidance. SG, YE, JM and DR designed the experiments. SG, KP, JM and JE conducted the experiments and analyzed the data. SG, YE and DR wrote the manuscript.

## Supplementary methods

### Reagents

Ethyl alcohol (190 proof) was purchased from VWR (Radnor, PA), and sucrose was purchased from Fisher Scientific (Pittsburgh, PA). The Qiazol RNA isolation kit was purchased from Qiagen (Redwood City, CA). cDNA was synthesized using the iScript cDNA Synthesis Kit (BioRad). Powerup SYBR Green PCR Master mix (ThermoFisher) was used for quantitative real-time PCR. Other common reagents were from Sigma Aldrich (St. Louis, MO) or Fisher Scientific (Pittsburgh, PA).

### Preparation of solutions

Alcohol solution was prepared from absolute anhydrous alcohol (190 proof) diluted to 20% alcohol (v/v) in tap water. Sucrose solution was diluted to 0.3% sucrose (v/v) in tap water.

### Collection of brain samples for biochemical analyses

Mice were euthanized 4 hours and 24 hours after the last drinking session (binge and withdrawal time point). Brains were removed and dissected on an ice-cold platform into 1 mm sections, and specific subregions-vlOFC, mOFC, and motor cortex (M2) were dissected based on Allen Brain Atlas.

### Quantitative real-time PCR

RNA was isolated using the RNeasy kit, and cDNA was synthesized using the iScript cDNA Synthesis Kit according to the manufacturer’s instructions. The resulting cDNA was used for quantitative real-time PCR, using Powerup SYBR Green PCR Master mix. Thermal cycling was performed on QuantStudio 5 real-time PCR System (Thermo Fisher Scientific Inc.) using a relative calibration curve. The quantity of *BDNF* mRNA was measured and expressed relative to *GAPDH* mRNA. PCR primers used: *BDNF* Forward 5’-TGC AGG GGC ATA GAC AAA AGG-3’, *BDNF* Reverse 5’-CTT ATG AAT CGC CAG CCA ATT CTC-3’, *GAPDH* Forward 5’-CGA CTT CAA CAG CAA CTC CCA CTC TTC C-3’ and *GAPDH* Reverse 5’-TGG GTG GTC CAG GGT TTC TTA CTC CTT-3’.

### Adeno-associated viruses

AAV2-DIO-Ef1a-BDNF-IRES-mCherry virus (AAV2-DIO-BDNF-mCherry; 3 x 10^12^ vg/ml) and AAV2-DIO-Ef1a-mCherry virus (AAV2-DIO-mCherry; 1 x 10^12^ vg/ml) were constructed in conjunction with C&M Biolabs (Richmond, CA) and were produced by the Duke University Viral Vector Core. AAV2-retrograde(retro)-Cre-GFP virus (AAVretro.hSyn.HI.eGFP-Cre.WPRE.SV40; 3x10^12^ vg/ml) was purchased from Addgene.

### Stereotaxic viral infection

Mice were anesthetized by vaporized isoflurane and placed on a digital stereotaxic frame (David Kopf Instruments). Two holes were drilled above the site of viral injection. The injectors (stainless tubing, 33 gauges; Small Parts Inc.) were slowly lowered into the target region. The injectors were connected to 10 µl Hamilton syringes, and the infusion was controlled by an automatic pump at a rate of 0.1 µl/min. The injectors remained in place for an additional 10 minutes to allow the virus to diffuse and then were slowly removed.

For circuit-specific expression of BDNF, mice received bilateral infusion of 1 µl of AAV2-DIO-Ef1a-BDNF-IRES-mCherry virus (AAV2-DIO-BDNF-mCherry, 3x10^12^ gc/ml) per hemisphere into the vlOFC (AP: +2.2, ML: ± 1.2, DV: -2.6) or M2 (AP: +2.2, ML: ± 1.3, DV: -1.35) and 1 µl of AAVretro-Cre-GFP (3x10^12^ vg/ml) into the DLS (AP: +1.1, ML: ± 1.9, DV: -2.95) or DMS (AP +1.1, ML +/- 1.2, DV -2.95). Control animals received 1 µl of empty vector: AAV2-DIO-Ef1a-mCherry virus (AAV-DIO-mCherry, 1x10^12^ gc/ml) in the vlOFC or M2 and 1 µl of AAVretro-Cre-GFP (3x10^12^vg/ml) in the DLS or DMS.

### Confirmation of viral expression

At the end of the experiments, animals were euthanized via cervical dislocation, the brains were removed, placed on ice, and dissected into 1 mm coronal sections. The fluorescent protein expressed by the virus (either GFP or mCherry) was visualized using an EVOS FL tabletop fluorescent microscope and images were obtained (ThermoFisher Scientific). Two animals that failed to exhibit fluorescence associated with viral overexpression were excluded from the study.

### Behavioral paradigms

#### Intermittent access to 0.3 % sucrose two-bottle choice

Sucrose intake paradigm was conducted as previously described (Hoisington et al., 2024). Animals received one bottle of 0.3% sucrose and one bottle of water 24 hours a day for two weeks on Monday, Wednesday, and Friday, with 24 or 48 hours (weekend) sucrose deprivation periods in which mice consumed only water. Sucrose solution intake (ml/kg), water intake (ml/kg), total fluid intake (ml/kg), and the preference ratio (volume of sucrose solution intake/total volume of fluid intake) were recorded every 24 hours for the duration of sucrose access. Corrections were made to account for spillage based on bottles affixed to an empty cage.

#### Open-field locomotion test

Mice were habituated to the room for 60 minutes prior to the experiment. Mice were placed in an open field apparatus (43cm x 43cm) in low-light conditions and allowed to explore for 10 minutes^51^. Locomotor activity was tracked using EthoVision XT software (Noldus, Leesburg, VA), and total movement (cm) and velocity (cm/s) were recorded. At the end of the session, the mouse was removed, and the apparatus was cleaned between sessions.

**Supplementary Table S1.**
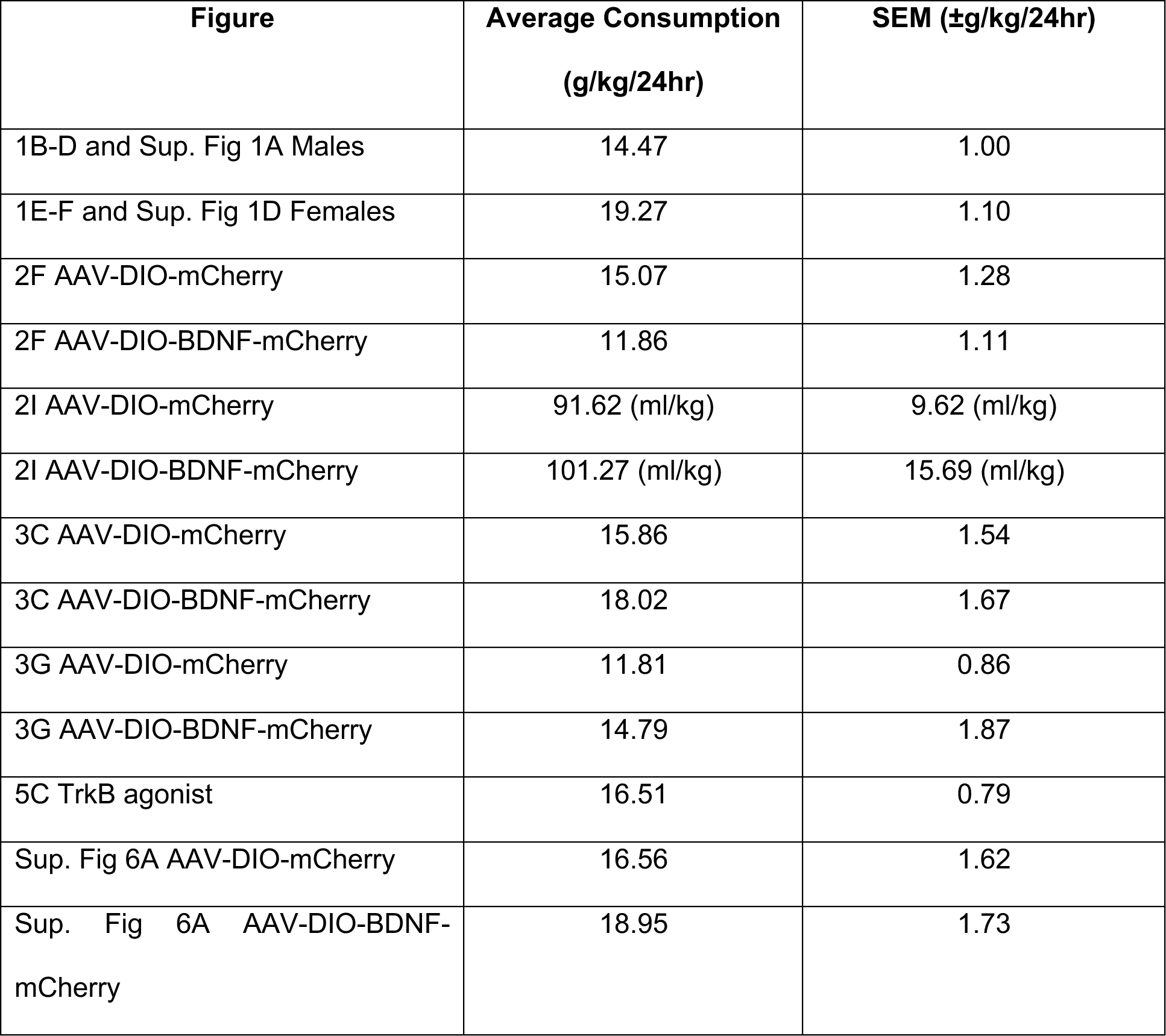
Average alcohol and sucrose consumption for behavior experiments. Average alcohol and sucrose consumption in mice subjected to the IA2BC with 20% alcohol and 0.3% sucrose, respectively.

### Supplementary Figures

**Supplementary Figure 1.**
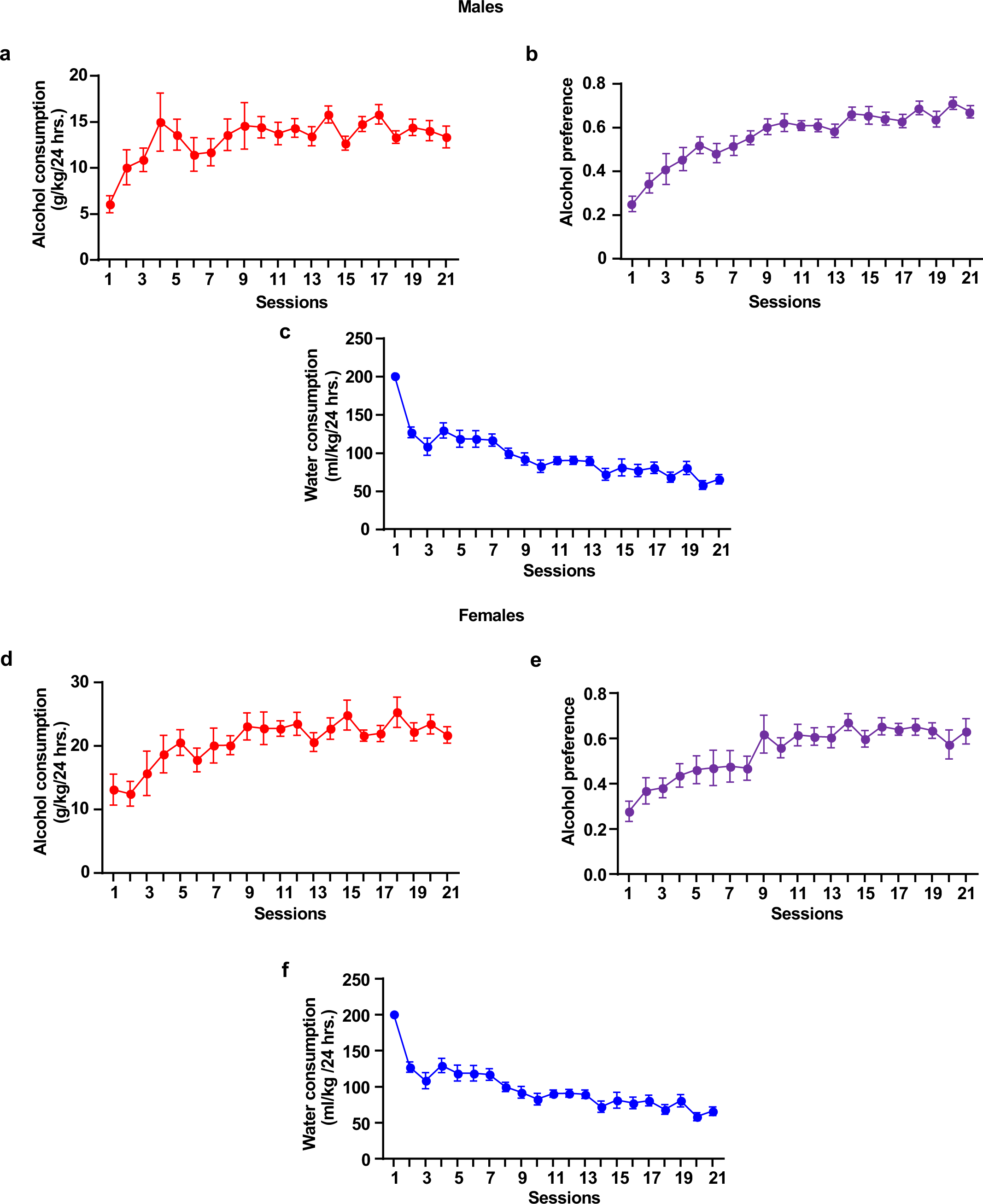
Drinking profile of male and female mice. Mice were subjected to IA20%2BC for 7 weeks in the home cage before harvesting vlOFC, mOFC and M2 regions for biochemical analysis. Alcohol consumption, preference, and water consumption of male (**a-c**) and female mice (**d-f**).

**Supplementary Figure 2.**
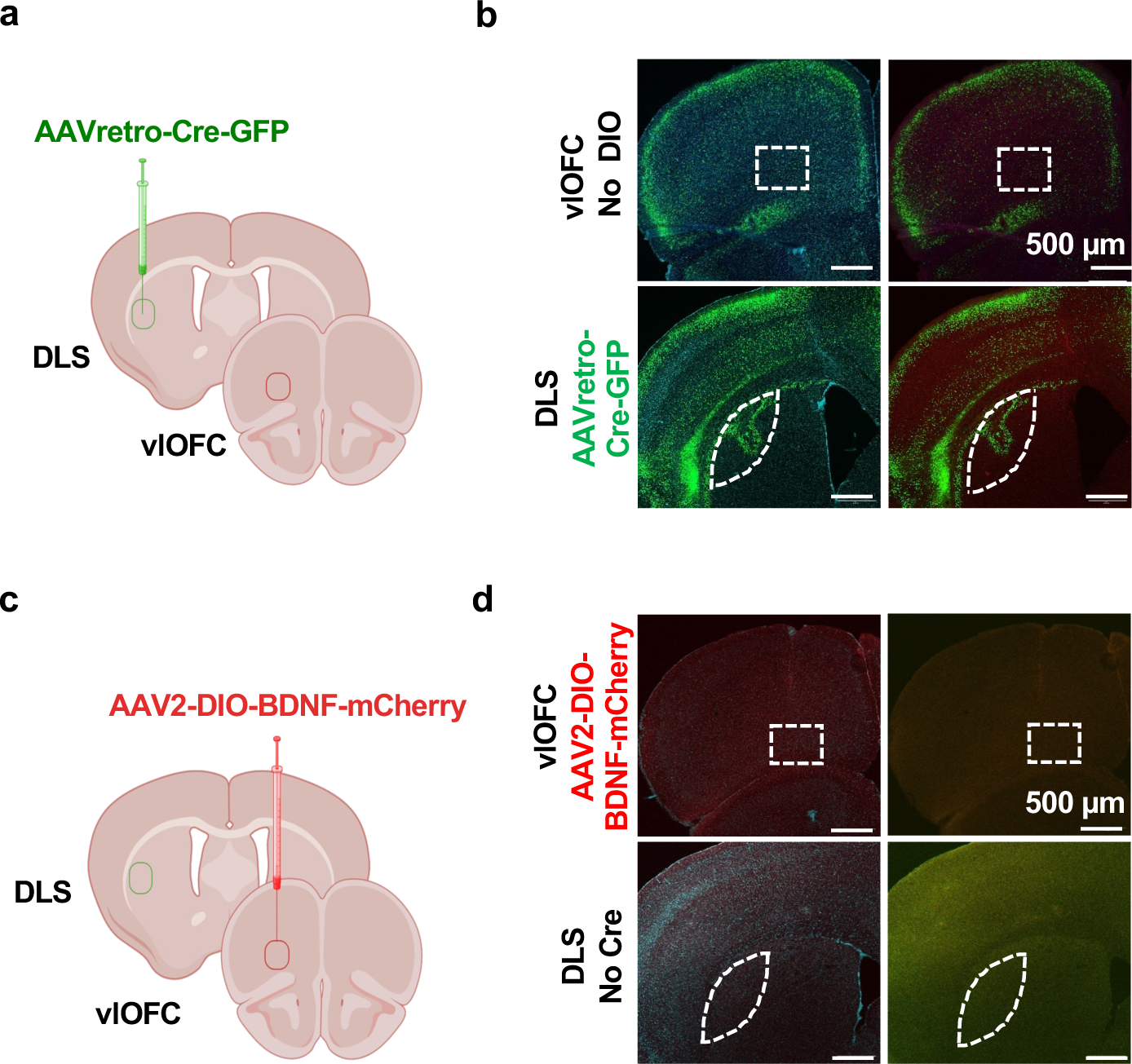
Confirmation of the Cre/DIO strategy efficacy. (**a**) Schematic representation of BDNF overexpression in vlOFC-to-DLS circuit. Mice received bilateral injections of AAVretro-Cre-GFP in the DLS. (**b**) Representative images of GFP expression (AAVretro-Cre-GFP) in the DLS and in cortical regions, including the vlOFC. (**c**) Mice received bilateral injections of AAV2-DIO-BDNF-mCherry in vlOFC. (**d**) Representative images showing no mCherry expression following AAV2-DIO-BDNF-mCherry injections in the vlOFC without AAVretro-Cre-GFP injection in the DLS

**Supplementary Figure 3.**
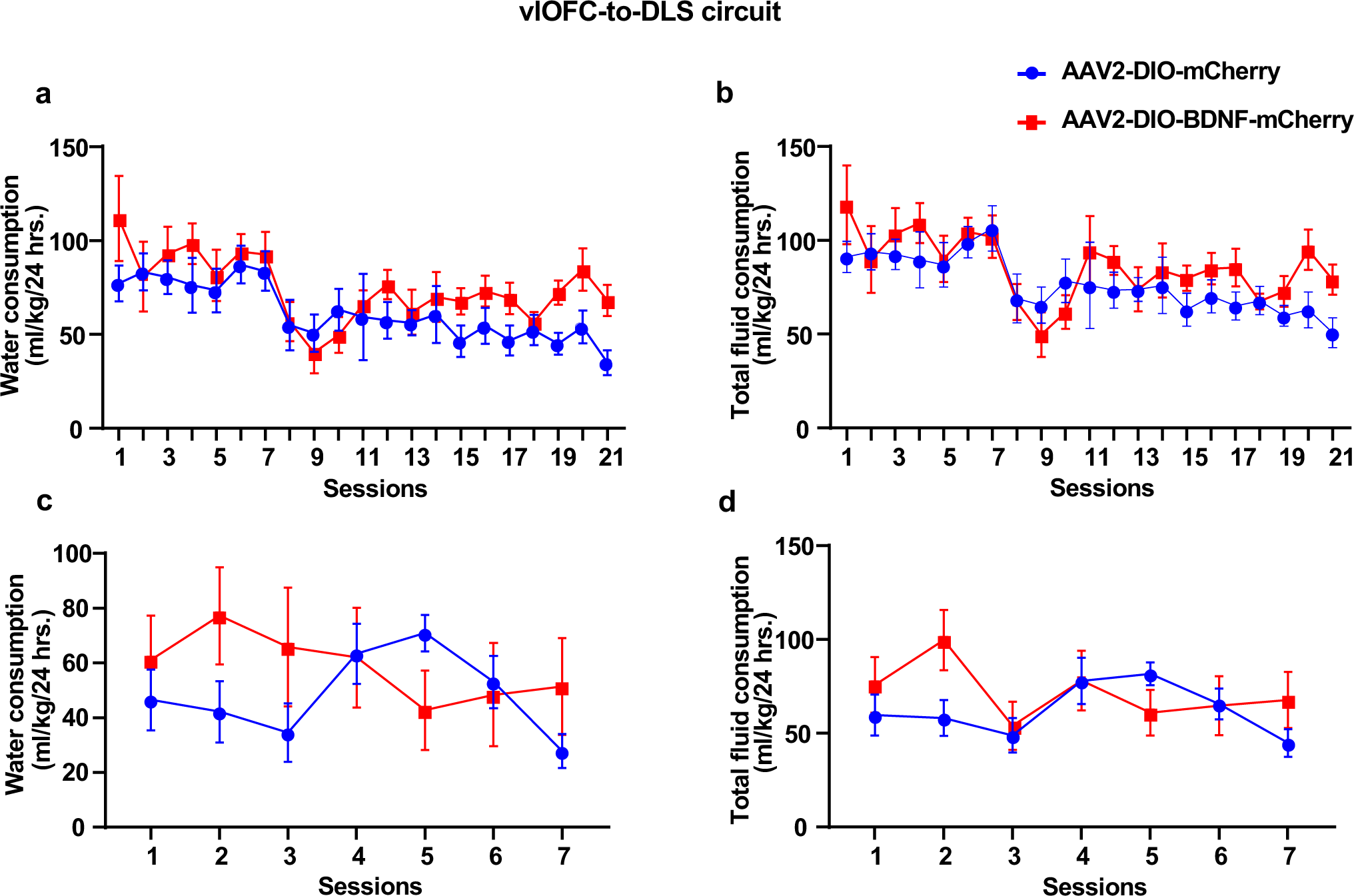
Overexpression of BDNF in vlOFC-to-DLS projecting neurons does not alter water or total fluid consumption during alcohol and sucrose intake. (**a-b**) Water and total consumption of alcohol was measured after BDNF overexpression in vlOFC-to-DLS projecting neurons **(a)** Water consumption was measured (Two-Way ANOVA, effect of BDNF overexpression, F _(1, 13)_ = 2.906, p =0.11, effect of session, F _(20, 245)_ = 4.437, ****p <0.0001, effect of interaction F_(20,245)_ = 0.92, p = 0.55). (**b**) Total fluid consumption was calculated (Two-Way ANOVA, effect of BDNF overexpression, F _(1, 13)_ = 1.888, p =0.19, effect of session, F _(20, 245)_ = 3.900, ****p <0.0001, effect of interaction F_(20,248)_ = 0.87, p = 0.61). (**c-d**) Water and total consumption of sucrose was measured after BDNF overexpression in vlOFC-to-DLS projecting neurons (**c**) Water consumption was measured (Two-Way ANOVA, effect of BDNF overexpression, F _(1, 11)_ = 0.2527, p =0.62, effect of session, F _(6, 58)_ = 0.8752, p =0.51, effect of interaction, F _(6, 58)_ = 2.479, p =0.03). (**d**) Total fluid consumption was calculated (Two-Way ANOVA, effect of BDNF overexpression, F _(1, 11)_ = 0.4471, p =0.51, effect of session, F _(6, 56)_ = 1.682, p =0.14, effect of interaction, F _(6, 56)_ = 2.386, p =0.04). Data are represented as mean ± SEM, n=6-9 per group

**Supplementary Figure 4.**
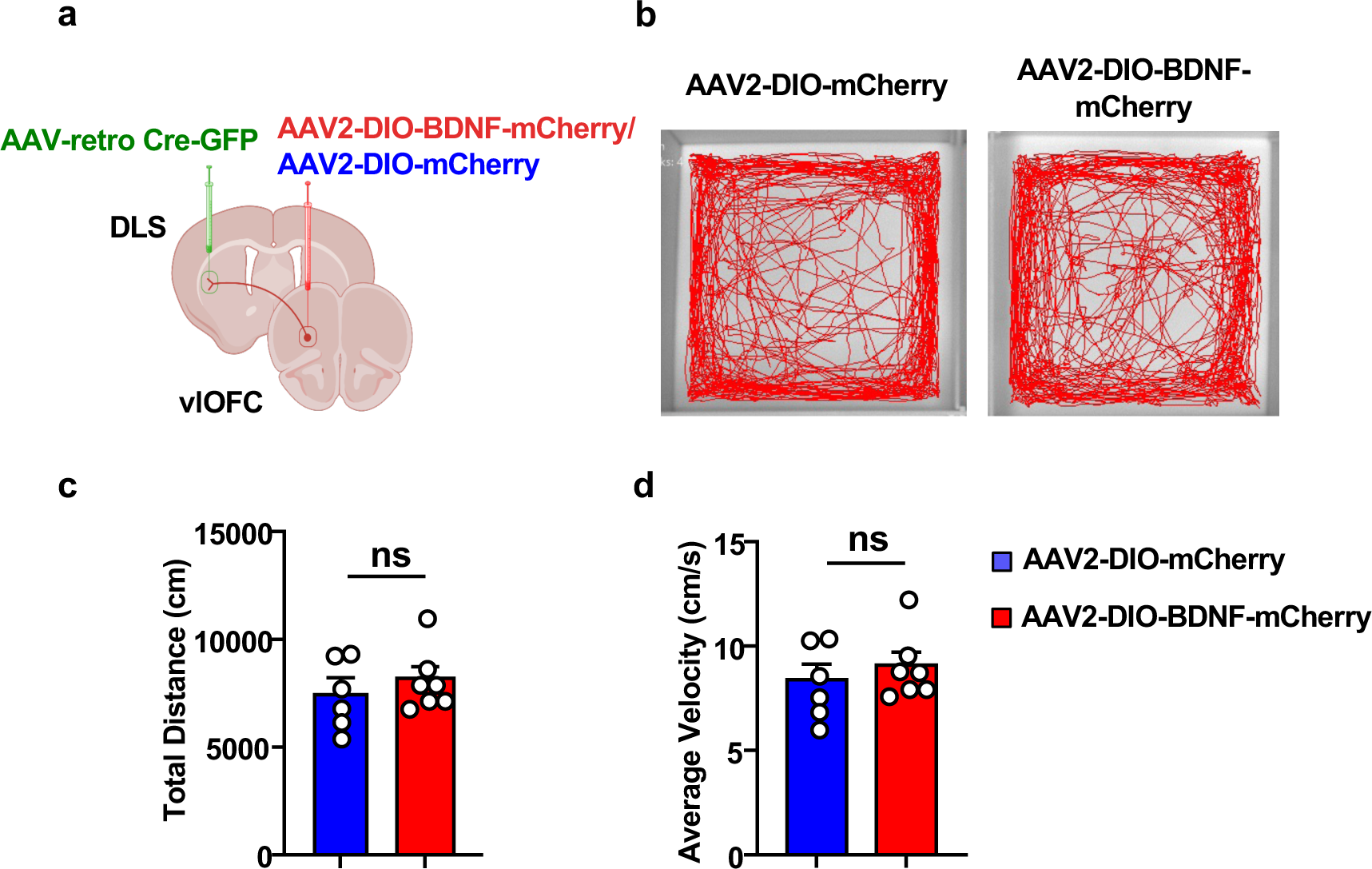
Overexpression of BDNF in vlOFC-to-DLS projecting neurons does not alter locomotion. (**a**) Experimental timeline: Mice received bilateral injections of AAV2-DIO-BDNF-mCherry in the vlOFC and AAVretro-Cre-GFP in the DLS. Three weeks after the surgery, mice were subjected to the open field test. (**b**) Representative tracks of the movement of AAV2-DIO-mCherry and AAV2-DIO-BDNF-mCherry mice. (**c-d**) Locomotion was recorded for 10 minutes, and total distance traveled (Mann-Whitney Test: U=20, p = 0.3969) and average velocity (Mann-Whitney Test: U=13, p = 0.9273) were calculated. Data are represented as mean ± SEM. ns: non-significant. n=6-7 per group.

**Supplementary Figure 5.**
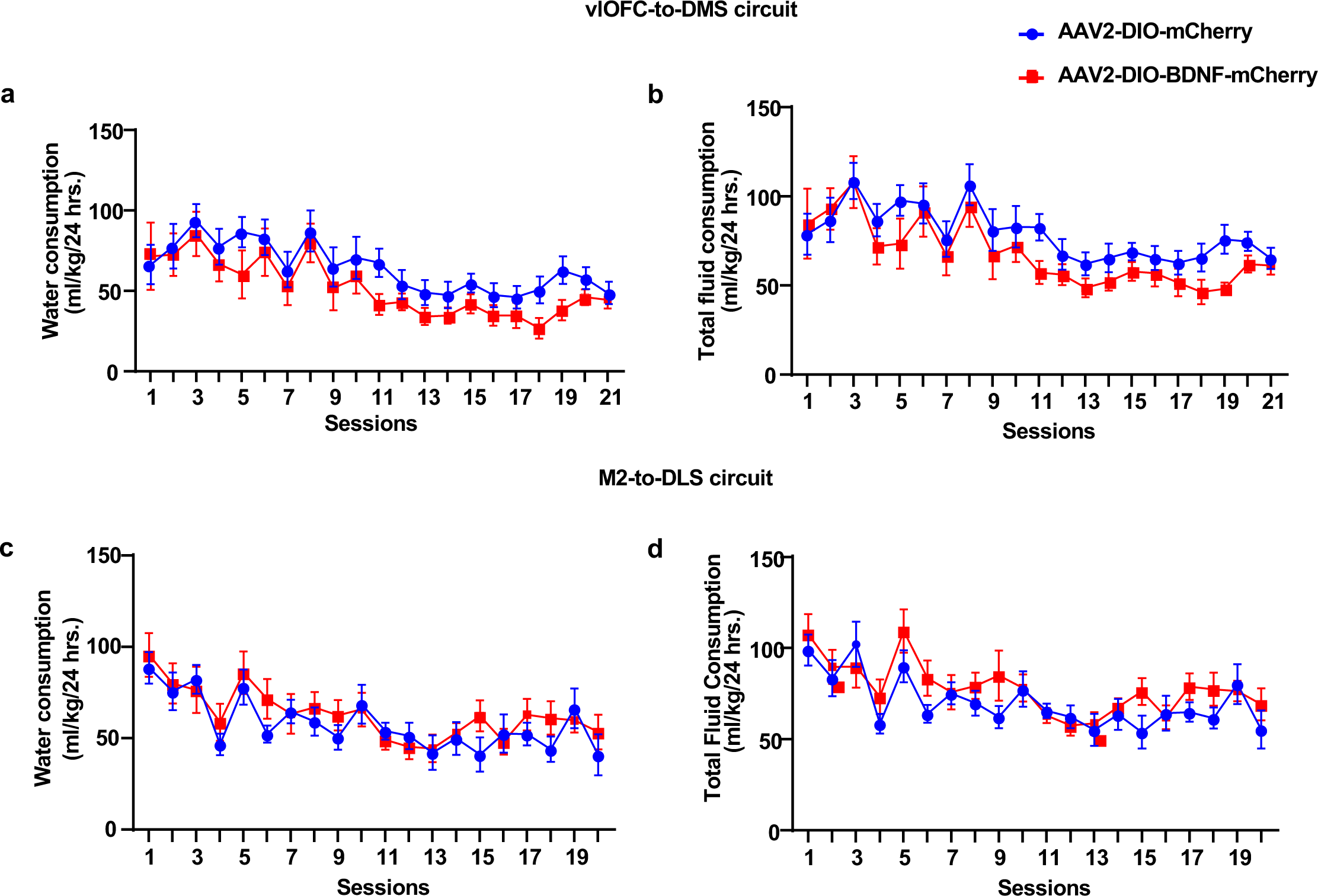
Overexpression of BDNF in vlOFC to DMS or M2 to DLS neurons does not alter water and total fluid consumption during voluntary alcohol intake in mice. (**a-b**) **vlOFC to DMS circuit. (a)** Water consumption was measured (Two-Way ANOVA, effect of BDNF overexpression, F _(1, 17)_ = 2.14, p =0.16, effect of session, F _(20, 329)_ = 7.296, ****p <0.0001, effect of interaction, F _(20, 329)_ = 0.38, p =0.99). (b) Total fluid consumption was calculated (Two-Way ANOVA, effect of BDNF overexpression, F (_1, 17_) = 2.042, p > 0.05, effect of session, F _(20, 325)_ = 7.631, ****p <0.0001, effect of interaction, F _(20, 325)_ = 0.63, p =0.88). (**c-d**) **M2 to DLS circuit.** (**c**) Water consumption was measured (Two-Way ANOVA, effect of BDNF overexpression, F _(1, 16)_ = 0.5373, p =0.47, effect of session, F _(19, 298)_ = 6.149, ****p <0.0001, effect of interaction, F _(19, 298)_ = 0.6853, p =0.83). (**d**) Total fluid consumption was calculated (Two-Way ANOVA, effect of BDNF overexpression, F _(1, 16)_ = 1.299, p =0.27, effect of session, F _(19, 300)_ = 6.18, ****p <0.0001, effect of interaction, F _(19, 300)_ = 0.92, p =0.55). Data are represented as mean ± SEM, n=8-10 per group

**Supplementary Figure 6.**
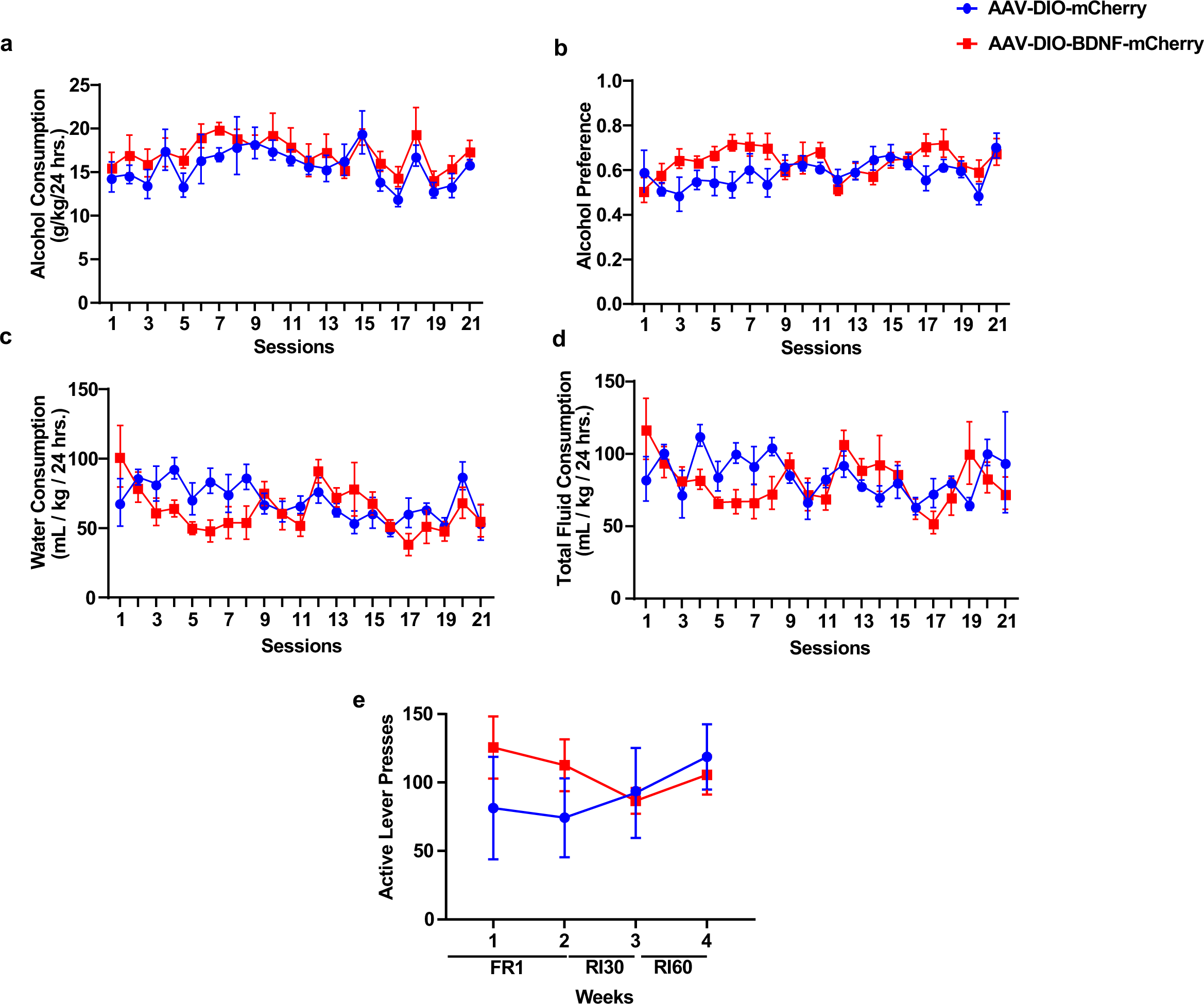
Drinking profile of mice prior to operant self-administration training and average lever presses of mice in operant training before undergoing surgery. Mice were subjected to IA20%2BC for 7 weeks in the home cage and were assigned to AAV-DIO-mCherry and AAV-DIO-BDNF-mCherry groups before they were trained to operantly self-administer alcohol. Alcohol consumption (**a**), preference (**b**), water consumption (**c**) and total consumption (**d**).(**e**) The group average of active lever presses during the FR1, RI30 and RI60 sessions in operant self-administration training before the animals undergoing the surgery (Two-way RM ANOVA: Effect of virus F_(1,13)_ = 0.249, p = 0.626; Effect of session F_(2.662, 34.61)_ = 1.931, p = 0.148; Effect of interaction F_(3,39)_ = 4.034, p = 0.013).

