## Supplementary material for "BDNF in Ventrolateral Orbitofrontal Cortex to Dorsolateral Striatum Circuit Moderates Alcohol Consumption and Gates Alcohol Habit"

| **Figure** | **Average Consumption (g/kg/24hr)** | **SEM (±g/kg/24hr)** |
| --- | --- | --- |
| 1B-D and Sup. Fig 1A Males | 14.47 | 1.00 |
| 1E-F and Sup. Fig 1D Females | 19.27 | 1.10 |
| 2F AAV-DIO-mCherry | 15.07 | 1.28 |
| 2F AAV-DIO-BDNF-mCherry | 11.86 | 1.11 |
| 2I AAV-DIO-mCherry | 91.62 (ml/kg) | 9.62 (ml/kg) |
| 2I AAV-DIO-BDNF-mCherry | 101.27 (ml/kg) | 15.69 (ml/kg) |
| 3C AAV-DIO-mCherry | 15.86 | 1.54 |
| 3C AAV-DIO-BDNF-mCherry | 18.02 | 1.67 |
| 3G AAV-DIO-mCherry | 11.81 | 0.86 |
| 3G AAV-DIO-BDNF-mCherry | 14.79 | 1.87 |
| 5C TrkB agonist | 16.51 | 0.79 |
| Sup. Fig 6A AAV-DIO-mCherry | 16.56 | 1.62 |
| Sup. Fig 6A AAV-DIO-BDNF-mCherry | 18.95 | 1.73 |

**Supplementary Table S1. Average alcohol and sucrose consumption for behavior experiments**

Average alcohol and sucrose consumption in mice subjected to the IA2BC with 20% alcohol and 0.3% sucrose, respectively.
